# High-resolution cryo-EM structure of integrin αIIbβ3 bound to disease-causing maternal HPA-1a antibody that blocks integrin activation

**DOI:** 10.64898/2026.03.06.709550

**Authors:** José M. de Pereda, Wendy Stam, Marcos Gragera, Femke van der Meer, Francisco J. Chichón, Eleftherios Zarkadas, Ellen van der Schoot, Gestur Vidarsson, Junichi Takagi, Coert Margadant

## Abstract

Integrins promote immunity, embryonic development, wound healing, and hemostasis, and are activated by ‘bent/closed’ to ‘extended/open’ conformational changes. Integrin αIIbβ3, being crucial for platelet activation and aggregation, is a therapeutic target for bleeding disorders and thrombosis. Human Platelet Antigen-1a (HPA-1a) on β3 is recognized by pregnancy-associated maternal alloantibodies, potentially causing fetal/neonatal alloimmune thrombocytopenia (FNAIT) and even intracranial hemorrhage or perinatal death. However, severe disease determinants are largely unknown. We report the first structure of an anti-HPA-1a antibody fragment (Fab 26.4) in complex with integrin αIIbβ3 at high resolution by cryo-electron microscopy. Fab 26.4 binding traps αIIbβ3 in the inactive, bent/closed conformation, is incompatible with integrin extension, and inhibits αIIbβ3-dependent fibrinogen binding and platelet aggregation. Thus, anti-HPA-1a antibodies directly impair integrin activation by preventing required conformational changes. These insights will improve FNAIT diagnostics and treatment, and spark the development of novel allosteric inhibitors against β3 integrins for future therapeutic applications.

## Introduction

The integrins constitute a family of 24 αβ heterodimeric transmembrane cell adhesion receptors that are crucial for immunity, embryonic development, wound healing, angiogenesis, and hemostasis (*1*). Dysregulated integrin function is involved in numerous pathological processes, and integrins are therefore therapeutic targets in a range of conditions including inflammation, multiple sclerosis, fibrosis, thrombosis, and cancer (2). While the integrin extracellular domains (ectodomains) bind soluble ligands in blood plasma, cell-surface proteins on other cells, or immobilized ligands in the extracellular matrix, their cytoplasmic tails connect to the cytoskeleton and recruit a variety of intracellular proteins, thus activating signaling pathways (*3*). The affinity of integrins for their ligands is regulated by conformational changes, most notably from the compact bent/closed conformation to the extended/open conformation, which is known as ‘integrin activation’ (*4*–*6*).

There are two integrins containing the β3-subunit; αvβ3, expressed on a variety of cells such as endothelial cells, and αIIbβ3 (also known as GPIIb/IIIa), which is expressed almost exclusively on megakaryocytes and platelets. Integrin αIIbβ3 is the most abundant glycoprotein on platelets, binds fibrinogen (FB) and von Willebrand Factor, and plays a crucial role in platelet aggregation and thus blood clotting (*7*). Reduced expression of αIIbβ3 or a failure in its activation causes inherited bleeding disorders such as Glanzmann’s thrombasthenia and leukocyte adhesion deficiency-III (*6, 8, 9*) Furthermore, αIIbβ3 is a target for alloimmune antibodies that can cause platelet destruction and bleeding, as occurring in the disorder post-transfusion purpura (PTP) in recipients of blood transfusion (*10*). In addition, alloimmune reactions can occur during pregnancy from mother to child, and constitute a major risk for pregnancy. Alloantibodies in pregnancy can be directed against paternally inherited antigens on fetal blood cells, including the human platelet antigens (HPAs) (*11*). This causes clearance of fetal platelets, leading to low platelet counts in the fetus and/or neonate, a condition called fetal/neonatal alloimmune thrombocytopenia (FNAIT) (*12*). Most severe FNAIT and PTP cases are caused by antibodies against HPA-1a on the integrin β3-subunit, which is due to a L33>P polymorphism occurring in 2.3% of the Western population (*13*). The clinical outcome of anti-HPA1a antibodies on the fetus is highly heterogeneous, ranging from asymptomatic to severe bleeding such as intracranial hemorrhage, which can result in lifelong neurological complications or perinatal death (*14*–*16*). The determinants of disease severity are to date largely unknown, which has hindered the development and implementation of prenatal diagnostic screening programs to identify women at high risk who would benefit from treatment (*17*).

We have recently found that anti-HPA-1a antibodies have a general preference for the bent/closed rather than the extended integrin conformation, and that their binding is decreased upon integrin conformational activation (*18*). Structural modeling of the epitope in different integrin conformations strongly suggests that HPA-1a is most accessible in the bent conformation, while in the extended integrin the epitope is partially occluded (*18*).

It is increasingly recognized that anti-HPA-1a antibodies are highly heterogeneous, both in epitope recognition and functional effects (*19*). While all antibodies recognize L33 on the β3 PSI domain, adjacent residues in the PSI, as well as the more distant I-EGF1 and I-EGF2 domains can contribute to antibody binding (*19*–*22*). Since the PSI/I-EGF regions are essential for integrin extension, anti-HPA-1a antibody binding may affect integrin activation (*23*), which is possibly related to disease severity in FNAIT (*19*). It is therefore crucial to identify the exact antibody interactions with HPA-1a, but structures of αIIbβ3 with disease-causing anti-HPA-1a antibodies do not currently exist.

Here we report the first cryo-electron microscopy (cryo-EM) structure of integrin αIIbβ3 in complex with the Fab fragment of a disease-causing HPA-1a antibody (Fab 26.4), which revealed high-resolution details of the anti-HPA-1a antibody-epitope interface. The structure suggests a mechanism for anti-HPA-1a antibody-mediated inhibition of integrin activation. Indeed, we find that Fab 26.4 inhibits αIIbβ3-dependent FB binding and platelet aggregation. The results will serve as a stepping stone for detailed analysis of how anti-HPA-1a antibody interactions with αIIbβ3 correlate with disease severity, which will have major impact on FNAIT research and diagnostic screening. In addition, we identify anti-HPA-1a antibodies here as allosteric inhibitors of β3 integrin function that could be used for therapeutic applications in the future, circumventing the adverse effects observed in the clinic of current antagonists of β3 integrins that target the ligand-binding site (*24*).

## Results

### Maternal anti-HPA-1a Fab 26.4 inhibits platelet integrin αIIbβ3 activation

The maternal anti-HPA-1a alloantibody 26.4 inhibits platelet aggregation (*25*), which could be due to effects of the antibody Fc tail or to direct effects of the Fab region on integrin function. To investigate this, we produced Fab 26.4 and tested its effects on integrin αIIbβ3-dependent FB binding and platelet aggregation. We first examined human FB binding induced by thrombin in platelets derived from four individual healthy donors (Fig. 1A). Fab 26.4 inhibited FB binding in all donors in a dose-dependent manner (Fig. 1B). In addition, Fab 26.4 dose-dependently inhibited platelet aggregation as determined by flow cytometry (Fig. 1C). This was further confirmed using the dual-color flow cytometry assay to test micro-aggregation (Fig. 1D-F), which we developed earlier (*26, 27*) to test platelet function using limited volumes of patient-derived samples with low platelet counts. This assay can therefore be used in the future to test functional effects of HPA1a antibodies on FNAIT case-derived samples.

**Figure 1.**
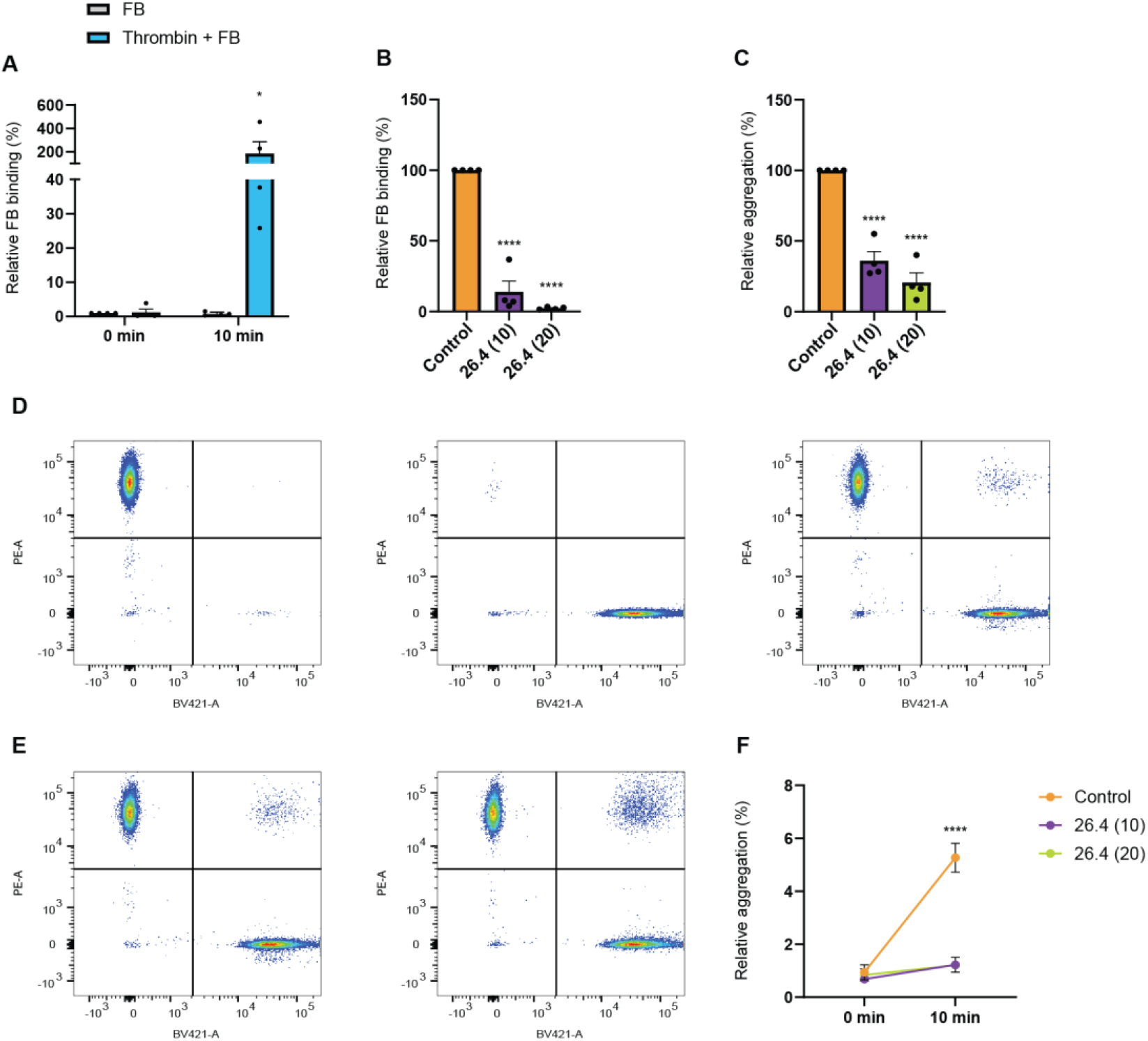
Anti-HPA-1a antibody Fab 26.4 inhibits integrin αIIbβ3-dependent FB binding and platelet aggregation. (**A**) Activation window of human platelets showing binding of FB (40 mg/ml), induced by thrombin (0.5 U/ml). Bars indicate means ± SEM of individual healthy platelet donors (n=4). (**B**) Human healthy donor platelets (n=4) were incubated with anti-HPA-1a antibody Fab 26.4 (10 or 20 µg/ml, 30 min) prior to thrombin-mediated activation, and binding of FB-650 (40 mg/ml) was measured by flow cytometry. Total platelets were gated based on forward/side scatter. Bars indicate means ± SD. (**C**) Thrombin/FB-mediated aggregation of human healthy donor platelets (n=4) is decreased after pre-incubation with Fab 26.4 (10 or 20 µg/ml, 30 min). Bars indicate means ± SD. (**D**) Gating strategy for the dual-color aggregation assay by flow cytometry. Healthy donor platelets, separately labeled with fluorescent tags, were mixed in a 1:1 ratio. Total platelets were divided into 4 quartiles based on fluorescent labeling with CellTrace Yellow (*left*), CellTrace Violet (*middle*), or the 1:1 mix of these populations (*right*). (**E**) Platelets were labeled as in (D), pre-incubated with Fab 26.4 (10 µg/ml, 30 min), and then exposed to thrombin (0.5 U/ml) and FB (40 mg/ml). Plots show the double-colored events, indicating aggregates, at 0 min (*left*) and 10 min (*right*). (**F**) Number of aggregates measured in the dual-color aggregation assay as outlined in (D,E), determined as the % double-colored events out of all colored events (Q2/(Q1+Q2+Q3)*100). Data points indicate means ± SD (n=2 donors). Statistical analysis was performed using two-way ANOVA with Šídák’s multiple comparisons correction (A,F), or Tukey’s multiple comparison correction (B,C). Statistically significant differences are indicated by asterisks (*p<0.05, ****p<0.0001).

Taken together, these findings demonstrate that the maternal, disease-causing antibody 26.4 blocks integrin αIIbβ3-dependent functions in platelets, in a manner dependent on its Fab region.

### Structure of integrin αIIbβ3 in complex with Fab 26.4

To understand how the 26.4 alloantibody regulates αIIbβ3, we determined the structure of the αIIbβ3/26.4 complex using cryo-EM. We produced soluble recombinant αIIbβ3 integrin ectodomains (αIIbβ3-ecto) using an extensively characterized strategy (*28, 29*). In these integrin constructs, the transmembrane and cytoplasmic segments are replaced by an inter-subunit disulfide-bridged clasp (Fig. 2A-B). We used Fab 26.4 instead of the full-length antibody because the Fab alone is sufficient to block αIIbβ3 activation (Fig. 1), while also avoiding the added complexity associated with antibody bivalency and the flexibility of the hinge that links the Fc to the Fab regions.

**Figure 2.**
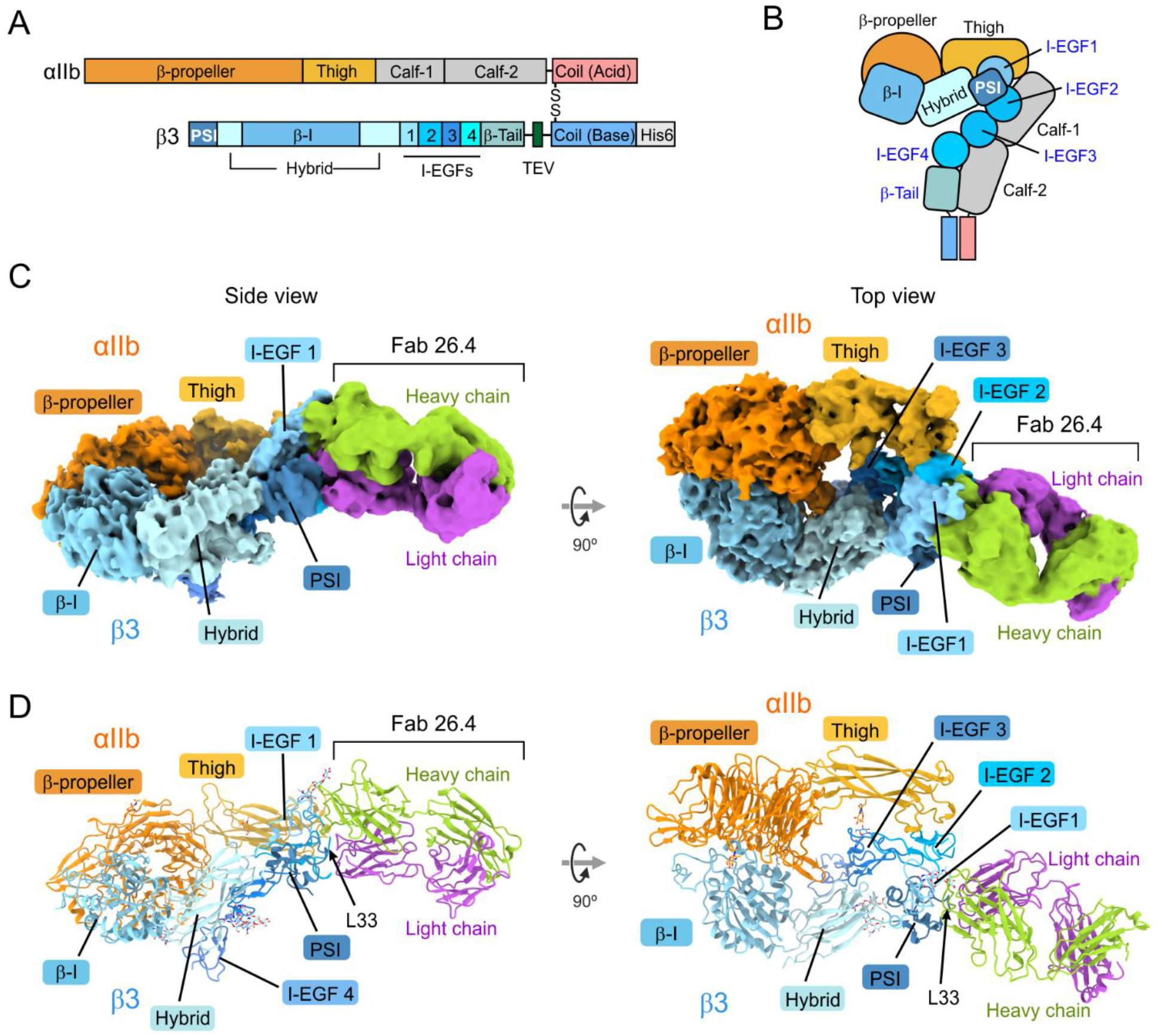
Cryo-EM structure of integrin αIIbβ3 in complex with Fab 26.4. (**A**) Domain organization of the primary sequences of the constructs that form the soluble αIIbβ3-ecto with a C-terminal covalent clasp. (**B**) Schematic representation of αIIbβ3-ecto in the bent/closed state. (**C**) Two orthogonal views of the cryo-EM map of the αIIbβ3-ecto in complex with Fab 26.4. Regions corresponding to domains in αIIb and β3 are colored as in (A). The Fab 26.4 heavy and light chains are shown each in a single color. (**D**) Ribbon representation of the atomic model of the complex, in the same orientations and colored as in (B).

Fab 26.4 formed a stable complex with αIIbβ3-ecto (fig. S1) that was suitable for structure solution using cryo-EM (fig. S2). The class featuring most of the αIIbβ3-ecto domain resulted in a map at an overall resolution of 2.64 Å (Fig. 2C). This map allowed the building of a model that included the β-propeller and thigh domains of αIIb, the PSI, hybrid, β-I and I-EGF1 to I-EGF4 domains of β3, and Fab 26.4 (Fig. 2D). No density was observed for the Calf-1 and Calf-2 domain of αIIb, which form the lower leg, and the C-tail domain of β3, probably because of high flexibility in these domains. The longest axes of the Fab and αIIbβ3 are approximately aligned, resulting in an elongated structure measuring ∼ 185 Å in length. Based on the orientation of αIIbβ3 with respect to the membrane revealed by recent cryo-EM structures of the full-length receptor, Fab 26.4 is oriented approaching the integrin from the membrane-proximal area (fig. S3).

### Details of the αIIbβ3/Fab 26.4 binding interface

The best-defined region in the overall cryo-EM map corresponds to the head of αIIbβ3, while the resolution was lower for the integrin legs and the Fab (fig. S2). The conserved domains of Fab 26.4 showed the lowest resolution, likely due to the inherent flexibility between the conserved and variable regions of Fab fragments. To enhance the resolution of the interface between αIIbβ3 and Fab 26.4, we performed local refinement, resulting in an improved map of the contact regions (Fig. 3A, Fig. 4 and fig. S2).

**Figure 3.**
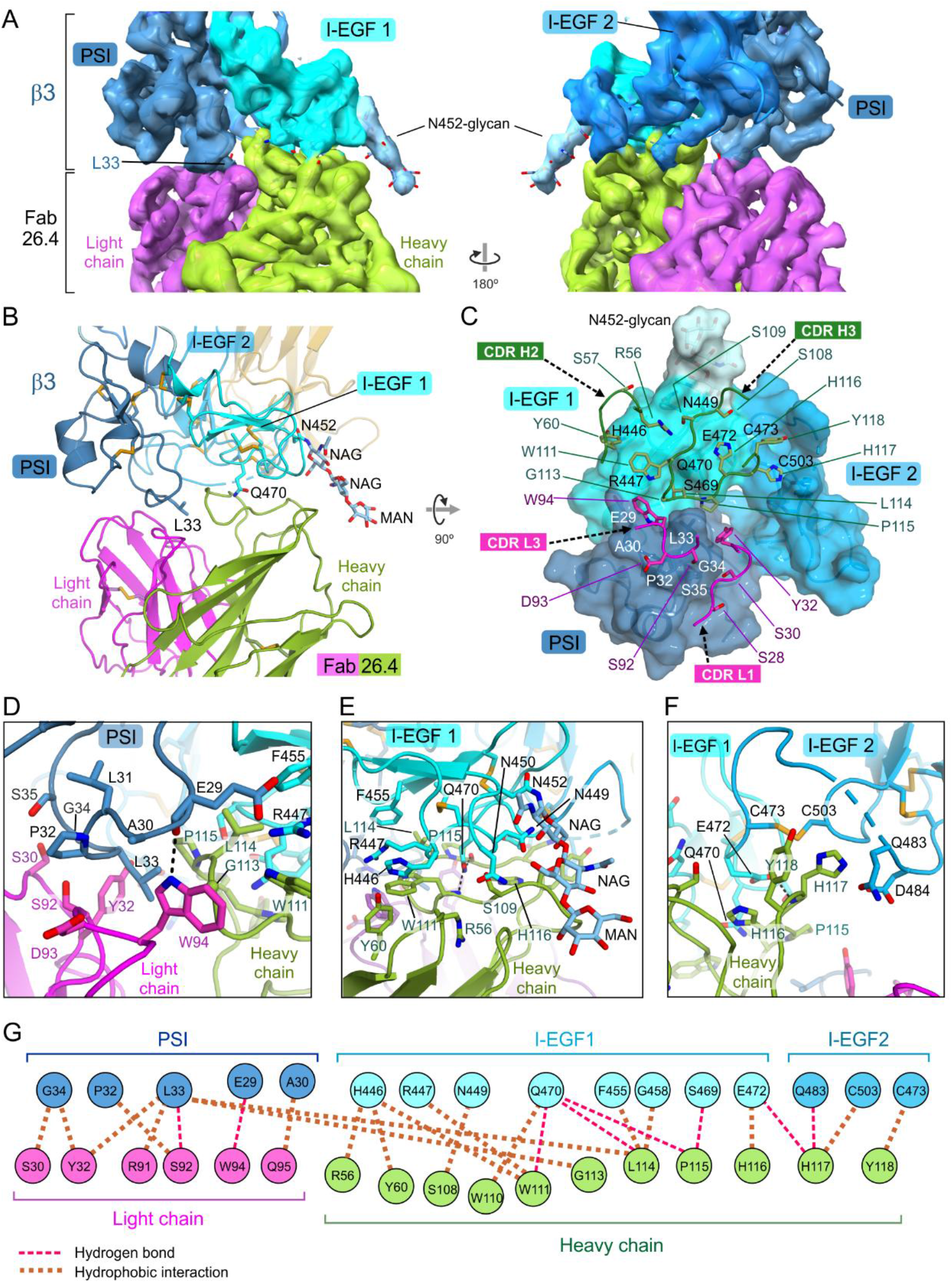
Interaction interface between Fab 26.4 and αIIbβ3. (**A**) Locally refined, unsharpened cryo-EM map showing the αIIbβ3/Fab 26.4 interface, with domains color-coded. (**B**) Ribbon diagram of the atomic model in the same orientation as (A, *left*), illustrating the interface architecture. (**C**) Footprint of Fab 26.4 on the β3-subunit. The β3 domains are shown in surface representation; Fab 26.4 CDR loop backbones contacting β3 are depicted as wires, and interacting side chains are shown as sticks. (**D**-**F**) Close-up views of contacts in three regions of the αIIbβ3/Fab 26.4 interface. Some hydrogen bonds between β3 and Fab 26.4, derived from the model, are shown as dashed lines. The N-linked glycan attached to N452 is also shown. NAG, N-acetyl-beta-D-glucosamine; MAN, alpha-D-mannopyranose. (**G**) Schematic diagram of the molecular contacts between β3 and Fab 26.4.

**Figure 4.**
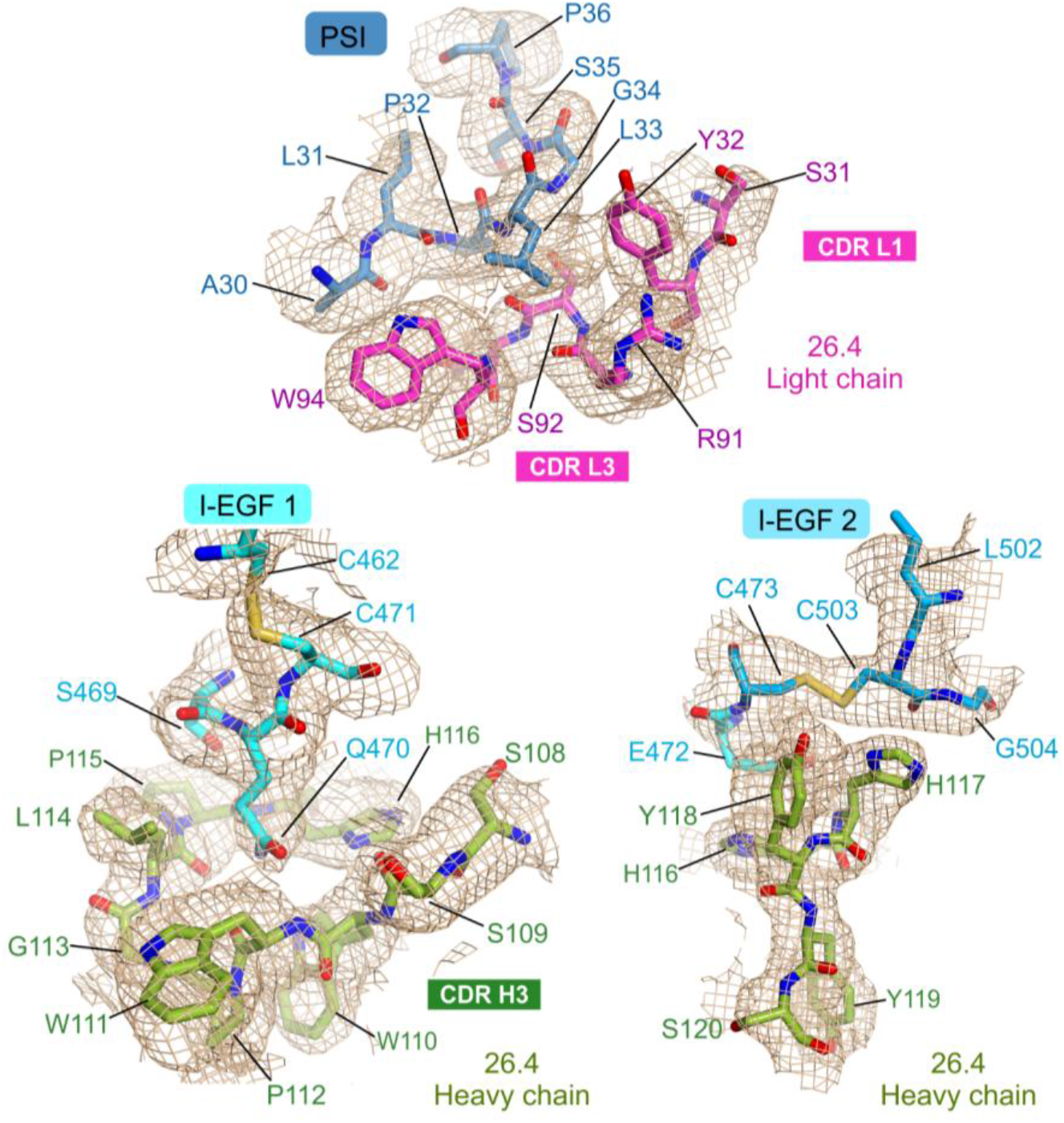
Details of the cryo-EM maps at the αIIbβ3/Fab 26.4 interface. Three regions of the sharpened cryo-EM density after local refinement focused on the β3/Fab 26.4 interacting domains. The regions of the refined atomic model fitted in the map are shown as sticks and color-coded by domain.

Fab 26.4 interacts exclusively with the β3-subunit, with two major contact areas (Fig. 3B,C). The light chain of 26.4 buries approximately 360 Å^2^ of the integrin’s solvent-accessible surface and primarily engages the β3 PSI domain, while the heavy chain buries an area of ∼750 Å^2^ and contacts predominantly the I-EGF1 domain.

The HPA-1a determinant, residue β3 L33, is sandwiched between the light and heavy chains of the antibody. Its side chain inserts into a pocket defined by residues Y32 and W94 from the complementarity-determining regions (CDR) L1 and L3 of the light chain, heavy chain (Fig. 3B-D, Fig. 4). PSI residues A30 and P32 also engage the Fab, consistent with previous observations (*20*). Noteworthy, D39 is not in or near the binding interface, although mutational analysis has previously implicated this residue in binding of 26.4 and other anti-HPA-1a antibodies (*20*). In contrast, several residues in the β3 PSI domain that have not been previously implicated in binding of anti-HPA-1a antibodies interact with Fab 26.4 or are close to the interface; these include E29, G34 and S35 (Fig. 3C-D, G).

The long CDR H3 of the Fab heavy chain (residues 105-118) engages the β3 I-EGF1 domain, primarily interacting with a surface centered on residues 469-472 (Fig. 3C). The side chain of β3 Q470 is surrounded by the CDR H3 and forms hydrogen bonds with backbone atoms of W111 and P115 in the Fab 26.4 heavy chain (Fig. 3E). Additionally, β3 residue E472 lies against H116 in the Fab (Fig. 3F). The heavy chain also interacts with the segment 446-460 of β3, with S108, W111, and L114 in the CDR H3 contacting N449, R447, and F455 in β3, respectively (Fig. 3E). In turn, W111 and L114 reduce the exposure of β3 C460 to the solvent. Other contacts include the interaction of Y60 in the CDR H2 with β3 H446. At the periphery of the I-EGF1/Fab 26.4 interface, the N-linked glycan chain attached to β3 N452 is oriented towards the Fab heavy chain; however, no specific interactions between the glycan moieties and the antibody are observed (Fig. 3A-C,E).

Importantly, Fab 26.4 also contacts the I-EGF2 domain. Residues Y118 and H116 in the heavy chain are within van der Waals distance of C473 and C503, which form the first disulfide bridge within the I-EGF2 domain (Fig. 3F and Fig. 4).

In summary, while β3 L33 is the antigenic determinant that triggers the generation of anti-HPA1a antibodies, 26.4 recognizes and engages a larger surface that extends throughout the PSI, I-EGF1 and I-EGF2 domains.

### Antibody 26.4 traps αIIbβ3 in an inactive conformation

In the αIIbβ3-ecto/Fab 26.4 complex, the integrin adopts a similar overall conformation as observed in the X-ray crystal structure of αIIbβ3-ecto (PDB: 3FCS) (*30*), and in the cryo-EM structures of full-length αIIbβ3 from human platelets in native lipid nanoparticles (PDB: 8T2V) (*31*), in detergent (PDB: 8GCD) (*32*), and in platelet membranes (PDB: 9E8A) (*33*) (Fig. 5A and fig. S4). The I-EGF domains 2, 3, and 4 of β3 are folded back towards the hybrid domain, in an arrangement characteristic of the bent inactive conformation (Fig. 5B). Furthermore, the head region formed by the αIIb β-propeller and the β3 β-I domain, which harbors the ligand-binding region, adopts an inactive conformation. This region superimposes perfectly onto the crystal structure of the inactive bent/closed state (3FCS) with a root mean square deviation (RMSD) of 0.787 Å (692 Cαs) (fig. S5A). The different activation states are characterized by the conformation of the three binding sites for divalent ions in the β-I region, which include a Mg^2+^ in the metal ion-dependent adhesion site (MIDAS) flanked by two Ca^2+^, one in the adjacent to MIDAS (ADMIDAS) site and the other in the synergistic metal ion-binding site (SyMBS) (*4, 34*). In addition, the positions of helices α1 and α7 are linked to the conformation of the metal binding sites. In our structure, the ion-binding sites and the adjacent helices α1 and α7 are in an inactive conformation (fig. S5B-D). In summary, αIIbβ3 in complex with Fab 26.4 is in an inactive, bent/closed conformation.

**Figure 5.**
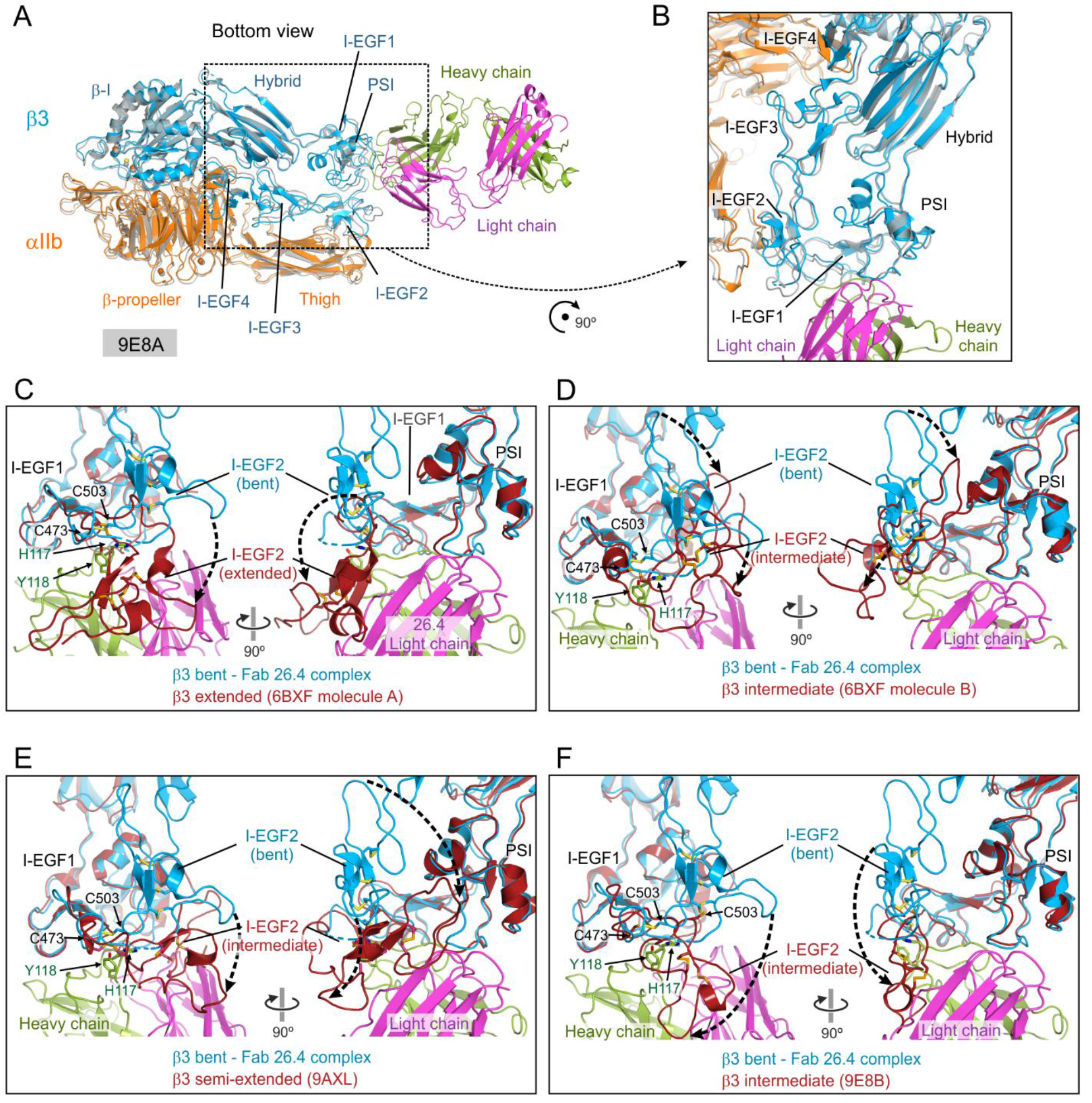
Comparison of the αIIbβ3/Fab 26.4 complex with activation-dependent αIIbβ3 conformations. (**A**) Ribbon representation of the complex (colored by protein chain) superimposed onto the cryo-EM model of full-length αIIbβ3 from human platelets in the bent, inactive conformation (PDB ID: 9E8A; shown in *gray*). For clarity, domains of the full-length structure not present in the complex are omitted. RMSD with 9E8A is 1.208 Å for 1190 Cαs atoms. (**B**) Close-up view of (A), highlighting the bent arrangement of the upper and lower legs of the β3-subunit. (**C**-**F**) Local superimpositions of the αIIbβ3/26.4 model (β3 in *blue*, Fab chains in *green* and *magenta*) with four β3 structures representing varying degrees of extension (*dark red-brown*). Dashed arrows indicate the apparent relative displacement of the I-EGF2 domain between bent and extended conformations.

Next, we analyzed if binding of Fab 26.4 would allow for conformational changes in the β3-subunit linked to integrin extension and activation. We compared the structure of our complex with structures of the β3-subunit in intermediate, partially extended, or fully extended states. First, we compared our structure to the structures of a monomeric β3 construct in extended (PDB: 6BXF molecule A) and intermediate (6BXF molecule B) conformations (*35*). After fitting the PSI and I-EGF1 domains, the I-EGF2 domain on the extended conformation is placed in a position occupied partially by Fab 26.4 (Fig. 5C). In the intermediate conformation of monomeric β3, the I-EGF2 domain also overlaps with the position of the Fab heavy chain (Fig. 5D). Next, we compared our structure with the semi-extended conformation of β3 trapped by the binding of Fab R21D10 to full-length αIIbβ3 (PDB: 9AXL) (*36*). The semi-extended I-EGF2 domain also partially collides with the position of Fab 26.4 (Fig. 5E). While the three previous intermediate or extended conformations are different from each other, they all relate to the bent conformation by movements that involve rotations around the disulfide bond between C473 and C503 in the I-EGF2 domain. These movements involve displacement of C503, which in the αIIbβ3-ecto/Fab 26.4 complex would be hampered by H117 from the Fab heavy chain, whose side chain contacts C503 (Fig. 3F and Fig. 5C-E). We also compared our complex to a recently described intermediate conformation observed by cryo-EM of full-length αIIbβ3 in human platelet membranes (PDB: 9E8B) (*33*) (Fig. 5F). In this state, the I-EGF2 would also collide with Fab 26.4. Noteworthy, the disulfide bridge between C473 and C503 is broken in this partially extended conformation.

Collectively, our observations suggest that Fab 26.4 traps αIIbβ3 in the bent state, and prevents the extension of β3 associated with integrin activation.

## Discussion

Here we report for the first time the structure of a disease-causing, maternal anti-HPA1a Fab fragment in complex with integrin αIIbβ3, solved by cryo-EM at high-resolution. The monoclonal antibody 26.4 investigated here is clinically relevant because it is derived from an alloimmunized mother who had an infant with severe thrombocytopenia and subcutaneous bleedings (*25*). Furthermore, a modified version of 26.4 was recently under consideration in a phase 2 clinical trial as a prophylactic approach against HPA-1a alloimmunization, but further development has been discontinued because of its poor pharmacokinetics in pregnant women (*37*). A deeper understanding of the biology of anti-HPA-1a antibodies is essential to identify determinants of severe disease in FNAIT, and for the development of diagnostic and therapeutic approaches.

Our structure clearly shows the interaction of Fab 26.4 with polymorphic residue L33, which is the basis for the alloimmune response leading to platelet destruction in HPA1a/b-incompatible pregnancies and platelet transfusions. The structure also reveals additional contacts in the PSI domain and extensive interactions in the neighboring I-EGF1 and I-EGF2 domains. We identify several residues in the β3 PSI and I-EGF1 domains that participate directly in the binding interface and that have been found previously in mutational studies as important for binding of anti-HPA1a antibodies (including 26.4), such as A30, P32, H446, and Q470 (*20*). Additionally, the structure reveals many novel β3 amino acids that are central to the interaction with the 26.4 antibody. In contrast, residues D39 and C460 are not directly at the interface, while it was found in earlier studies that mutation of these residues disrupted the binding of 26.4 and/or other anti-HPA-1a antibodies (*20*). C460 sits at the bottom of a pocket (∼ 3.9 Å from the side chain of L114 in the Fab heavy chain), and is therefore not in direct contact with the Fab. While not directly involved in anti-HPA-1a antibody binding, mutation of D39 and/or C460 probably induces local conformational changes that disrupt the HPA-1a epitope (and in the case of C460, the disulfide bond that is formed by this residue). We also observe unequivocal density for the N-linked glycan at N452 in the β3 I-EGF1 domain. Although no specific interactions with the Fab are observed, its proximity to the Fab heavy chain suggests that this glycan might contribute to antibody recognition, and that changes in the glycan composition could affect binding of 26.4 to β3.

The integrin in our complex is clearly in bent/closed conformation. The I-EGF domains 2 to 4 in the β3 lower leg are folded back towards the hybrid domain. In consonance with the bent conformation, the ligand-binding site in the αIIbβ3 head region (formed by the αIIb β-propeller and the β3 β-I domain) adopts an inactive state. Our structure showed unequivocal density for helices α1 and α7 in the β3 β-I domain, whose conformation is linked to the activation state of the adjacent ion-binding sites, SyMBS, MIDAS and ADMIDAS (*4, 34*). We and others had previously shown that anti-HPA-1a antibodies have a general preference for bent/closed integrins rather than extended (active) integrins (*18, 23*). Comparison of our structure with structures of αIIbβ3 in intermediate or fully extended conformation now reveals that binding of Fab 26.4 clashes with the I-EGF2 upon extension, offering a mechanistic explanation for those observations. Furthermore, our analysis predicts that Fab 26.4 binding will prevent or inhibit αIIbβ3 (and probably also αvβ3) integrin conformational activation, by impeding I-EGF1/2 domain extension along the hinge region. In addition to causing steric hindrance with the I-EGF2 domain upon extension, the Fab contacts the disulfide bond between C473 and C503. This prevents any movement around this Cysteine pair and may counteract disulfide bond disruption, which is linked to αIIbβ3 extension and activation (*38*). Therefore, Fab 26.4 (and possibly other anti-HPA-1a antibodies) might block the early stages of β3 extension leading to prevention of integrin activation. Indeed, we find that Fab 26.4 prevents fibrinogen binding to platelets and platelet aggregation, two processes dependent on αIIbβ3 activation. Our results may explain previous observations of anti-HPA-1a antibodies blocking several β3 integrin-dependent processes on platelets, endothelial cells, and trophoblasts (*19, 25, 39*–*44*), and suggest that this is not due to effects of the Fc tail, such as steric hindrance or binding to Fc receptors that may trigger downstream signaling responses that impair integrin activation. The exact orientation and binding of anti-HPA-1a antibodies is probably of utmost importance for their functional effects on integrin activation. In this respect, it is noteworthy that the monoclonal antibody R21D10, which is not an HPA-1a-specific antibody but it does bind the PSI domain including L33, allows for partial extension of the I-EGF1/I-EGF2 segment. R21D10 stabilizes this ‘semi-extended’ conformation and blocks ligand binding by steric hindrance (*36*).

While our structure reports the interaction of αIIbβ3 with a function-blocking anti-HPA-1a antibody, it is expected that there is considerable heterogeneity in the structural and functional features of anti-HPA-1a antibodies. Furthermore, we used here the purified αIIbβ3 ectodomain, rather than the full-length integrin embedded in a lipid environment, as occurs at the cell-surface. Nevertheless, our structure constitutes an important starting point for future work including cryo-EM studies aimed at elucidating the exact differences between HPA-1a antibodies that cause severe disease and others that only induce mild defects. Such studies are ongoing and are expected to generate a wealth of information with a broad range of applications, from a diagnostic test that can predict pregnancies at high-risk to develop major complications caused by anti-HPA-1a alloantibodies, to the design of modified therapeutic antibodies, aimed at preventing alloimmunization. In addition, a thorough understanding of anti-HPA-1a antibody interactions and functional properties will also generate novel opportunities for the development of allosteric β3 integrin inhibitors that target the PSI/I-EGF1/2 regions, rather than the commonly targeted ligand-binding site. Such inhibitors may work as pure antagonists, in contrast to many previously developed inhibitors that unexpectedly acted as partial agonists and therefore have caused major side effects in the clinic (*2, 24*).

## Materials and methods

### Experimental design

The objective of the study was to understand the functional and structural interactions of human anti-HPA-1a alloantibodies with integrin αIIbβ3. To this end, we produced a complex of the Fab fragment of antibody 26.4 (derived from a mother who had an infant with thrombocytopenia and bleedings), with the purified integrin αIIbβ3 ectodomain. We collected and analyzed cryo-EM datasets of the complex and compared these to other existing αIIbβ3 structures in different conformations. In addition, we determined the functional effects of Fab 26.4 on platelets from human donors.

### Platelet isolation, activation, and aggregation assays

Human platelets were isolated from healthy anonymous volunteers. Citrated whole blood was centrifuged at 150g for 20 min at room temperature without brake. The platelet-rich plasma was harvested and supplemented with 300 nM prostaglandin E1 (PGE1, Merck) before centrifugation at 480g for 15 min without brake to pellet platelets. Platelets were resuspended in warm PIPES/saline/glucose (PSG) buffer (5 mM PIPES, 45 mM NaCl, 4 mM KCl, 50 μM Na_2_HPO_4_, 1 mM MgCl_2_, and 5.5 mM D-glucose, pH 6.8) containing 300 nM PGE1, followed by centrifugation at 480g for 15 min. The washed platelets were resuspended in PSG/1% BSA and counted (Casy counter).

To examine integrin αIIbβ3 activation, platelets (2×10^6^/condition) were incubated with or without Fab 26.4 (10-20 µg/ml) in PSG/1% BSA without PGE1 for 30 min. Next, platelets were stimulated with thrombin (0.5 U/ml) and 40 µg/ml fluorescently labeled human FB purified from plasma (Haemacomplettan P, purchased from CSL Behring) for 10 min, followed by fixation with 1% PFA. FB was labeled in-house with DyLight650 (Thermo Scientific, ThermoFisher) according to the manufacturers’ instructions. FB bound to platelets was measured by flow cytometry on an LSRFortessa or LSRII machine (BD biosciences) and analyzed using FlowJo software (v10.9).

To test platelet aggregation, platelets were labeled with either CellTrace Yellow (2.5 µM, Invitrogen) or CellTrace Violet (5 µM, Invitrogen) and mixed in a 1:1 ratio. Labeling occurred for 20 min at 37°C in PSG buffer with PGE1, followed by 5 min incubation in 2% BSA/PSG buffer with PGE1 and centrifugation twice for 7 min at 800g. Labeled platelets were subsequently incubated with or without Fab 26.4 (10-20 µg/ml) in PSG/1% BSA for 30 min prior to stimulation with thrombin (0.5 U/ml) and FB-DyLight650, and fixation. Aggregation was measured directly and after 10 min by flow cytometry, using FSC/SSC scatter, as well as the dual-color micro-aggregation assay as described previously (*26, 27*).

### Protein expression and purification

The human αIIbβ3-ectodomain (αIIbβ3-ecto) was expressed in CHO-lec 3.2.8.1 cells using the following two constructs. One construct coded for the αIIb residues 1-963 followed by a site recognized by the tobacco etch virus (TEV) protease, and an acidic α-helical coiled-coil, in the pEF1/V5-HisA vector with neomycin resistance (*30*). The other construct coded for the β3 residues 1-692, followed by a TEV site, a basic α-helical coiled-coil, and a hexahistidine tag in the pEF1-Puro vector with puromycin resistance (*28*). The C-terminal acidic and basic α-helices in αIIb and β3 form a coiled coil clasp that is further stabilized by an inter-chain disulfide bridge (*28*–*30*). Secreted αIIbβ3-ecto was purified from the culture media as described earlier (*28*). Briefly, proteins in the cell culture supernatant were concentrated by ammonium persulfate precipitation, and subsequently resuspended in 20 mM Tris-HCl, 1 M NaCl, 20 mM imidazole, 1 mM CaCl_2_, 1 mM MgCl_2_ (pH 8.0). Next, αIIbβ3-ecto was purified by immobilized metal chelate affinity chromatography (IMAC) using a 5 ml His-Trap HP column (Cytiva), followed by elution with 20 mM Tris-HCl, 200 mM NaCl, 250 mM imidazole, 1 mM CaCl_2_, and 1 mM MgCl_2_, (pH 8.0). Subsequently, the protein was subjected to size exclusion chromatography on a Superdex 200 10/300 column (Cytiva), equilibrated in buffer A (50 mM Tris-HCl, 150 mM NaCl, 1 mM CaCl_2_, 1 mM MgCl_2_, pH 7.5). Then, αIIbβ3-ecto was subjected to ion exchange chromatography on a MonoQ 4.6/100 PE column (Cytiva). Finally, the integrin was extensively dialyzed against buffer A, and concentrated by ultrafiltration using Amicon Ultra devices to ∼1 mg/ml αIIbβ3-ecto. The human maternal monoclonal antibody 26.4 against HPA-1a has been originally cloned from B-cells of an HPA-1a-alloimmunized woman who had an infant with FNAIT (*25, 44*). Recombinant Fab 26.4 was produced in HEK-FS cells and purified as described previously (*45*).

### In vitro reconstitution and purification of the αIIbβ3/Fab 26.4 complex

The complex was formed by mixing ∼4.5 µM αIIbβ3-ecto with 1.1 to 1.6 molar excess of Fab 26.4, and was purified by size exclusion chromatography on a Superdex 200 10/300 GL column (Cytiva) equilibrated in buffer A; 100 µl of the mixture was injected in the column. The complex eluted as a single peak (fig. S1) and was concentrated by ultrafiltration to ∼1.3 mg/ml. Complex formation was confirmed by analysis of the samples by SDS-PAGE (8% acrylamide gel) under non-reducing conditions.

### Cryo-EM sample preparation and data collection

Grids were prepared at the Cryo-EM Facility of the CNB-CSIC (Madrid). Grids were glow-discharged for 30 sec at 25 mA using a EMITECH K100X device. Vitrification was performed using a Vitrobot Mark IV (Thermo Fisher Scientific) at 4°C and 95% humidity. 3 µl of purified integrin αIIbβ3-ecto/Fab26.4 complex at 0.13 mg/ml was applied to Quantifoil Cu/Rh R 0.6/1 grids (Quantifoil Micro Tools GmbH, blotting time 3 sec, blot force 3), followed by plunge freezing into liquid ethane. Initial grid screening was carried out on a Talos Arctica (Thermo Fisher Scientific) at CNB-CSIC (Madrid), operating at 200 kV and equipped with a Falcon 4i direct electron detector. The grids showed even distribution of particles, and the obtained 2D-3D data confirmed the quality of the sample to continue with acquisition of a large dataset at a high-end microscope.

Final data used for structure solution were collected on a Titan Krios G4 (Thermo Fisher Scientific), equipped with a cold field emission gun, Selectris X energy filter (10 eV slit width), a Cs of 2.7 mm, and a Falcon 4i direct electron detector. Data acquisition was performed at the CM02 beamline of the ESRF synchrotron, the French CRG cryo-EM beamline operated by IBS–ISBG.

Images were recorded at 165,000× nominal magnification, corresponding to a pixel size of 0.73 Å. The data were acquired in counting mode, with 50 fractions generated from 100 raw frames per movie (∼0.8314 e^−^/Å^2^/fraction). The total exposure time was 2.93 s, corresponding to a total dose of 41.57 e^−^/Å^2^, delivered at 8.21 e^−^/pixel/s. The defocus range was -0.5 to -2.2 µm, in 0.1 µm increments. Three images per hole were acquired. Image acquisition parameters and data processing statistics are summarized in table S1. Raw image stacks were saved in TIFF format and were not gain-corrected.

### Cryo-EM image processing

Single Particle Analysis was carried out within the Scipion platform (*46*). A total of 41,016 movies were aligned, dose-weighted, and initially downsampled to a pixel size of 1.46 Å/px with MotionCorr (*47*). Next, a sequence of quality filters was applied for micrograph curation: (I) xmipp s– movie maxshift (*48*), to discard movies with maximum frame-to-frame shift or global movie shift greater than 10 or 30 Å, respectively, (II) xmipp – tilt analysis (*48*), to exclude tilted micrographs based on analyzing power spectral density correlations across image quadrants and, finally, (III) miffi (*49*), keeping only micrographs labeled as “Good”. The number of micrographs for subsequent analysis was 25,980. Contrast Transfer Function (CTF) was estimated with ctffind (*50*), retaining micrographs with resolution better than 6 Å in a defocus range between 1,000-40,000 Å. Particle picking was performed with crYOLO (*51*) on low-pass filtered micrographs with automatic estimation of the box size. 6,717,934 particles were extracted at 2.42 Å/px and 128 box size and subjected to three rounds of 2D classification with cryoSPARC (*52*). The resulting particles (3,027,507) were used to generate 6 initial models with *ab initio* reconstruction in cryoSPARC, followed by 3D Heterogeneous Refinement. From here, particles for the class featuring the ternary complex were selected, reextracted at 1.46 Å/px and 212 box size and subjected to Non-uniform refinement (*53*), reaching Nyquist resolution of 3 Å. Next, the subset of particles was further cleaned with the 3D classification protocol in cryoSPARC, without recalculating alignments, into 8 classes (initialization mode: PCA, target resolution: 7 Å). Particles from the selected class (84,349) were reextracted again from non-downsampled micrographs using a 424-px box, Fourier-cropped to 300 px, resulting in a pixel size of 1.03 Å/px. A final refinement using these particles reached 2.6 Å resolution. The cryo-EM processing flow-chart is summarized in fig. S2.

For local refinement of the epitope-paratope interface, particles were re-centered on the region of interest, re-extracted, and assigned the angles from the previous global refinement. The reference volume was obtained after homogeneous reconstruction in cryoSPARC, followed by local refinement with rotation search extent of 40 degrees and shift search extent of 10. The mask for local refinement included the variable regions of Fab 26.4 and the PSI and I-EGF1/2 domains of the β3-subunit of the complex.

Maps were sharpened with deepEMhancer (*54*), and resolution was reported according to the Gold Standard Fourier Shell Correlation (GS-FSC) method, considering 0.143 as threshold. Local resolution was calculated with phenix.local_resolution (*55*).

### Model building and refinement

The model of αIIbβ3-ecto in the complex was built starting from the crystal structure of a similar construct in the bent/closed conformation (PDB: 3FCS) (*30*). Initial docking of the model onto the full cryo-EM map (unsharpened) was done interactively with Coot (*56*). The αIIb-subunit was positioned by rigid-body real space fitting the β-propeller followed by fitting of the thigh domain. Similarly, the β3-subunit was placed by first fitting the β-I domain on the map, and subsequently the Hybrid, PSI, and I-EGF1-4 domains. Ca^2+^ and Mg^2+^ cations and coordinating H_2_O molecules were initially modeled as in the 3FCS structure. Modeling of Fab 26.4, which consists of the variable and conserved domains of the light and heavy chains, was done using a custom-made model predicted by AlphaFold3 (*57*). The Fab model was first fitted as a single rigid body onto the map and subsequently each of the four domains were individually fitted.

The model of the complex was refined by iterative cycles alternating manual modeling in Coot using the full and local refinement cryo-EM maps (unsharpened and sharpened), with real-space refinement against the full unsharpened map using Phenix (*58*). Distance restraints for the metal coordination’s, and secondary structure and torsion restraints derived from the αIIbβ3 crystal structure were used during refinement. The quality of the structures was evaluated against the unsharpened full map using Phenix validation tools (*58*) and MolProbity (*59*). The final model includes the β-propeller and thigh domains of αIIb (residues 1-599), the PSI, hybrid, β-I and I-EGF domains 1 to 4 of β3 (residues 1-74, 79-478, and 483-600), the heavy chain (residues 1-238), and the light chain (residues 1-214) of Fab 26.4. N-linked glycans were modeled attached to αIIb residues N15 and N570, and to β3 residues N99, N320, N371, N452, and N559. The statistics of the refined structure are shown in table S1.

Molecular figures were created with UCSF ChimeraX (v1.10) (*60*) and Pymol (*61*). The αIIbβ3/Fab 26.4 interface was analyzed with PISA (*62*) and DIMPLOT-LigPlot+ (*63*).

### Statistical analysis

Flow cytometry data for FB binding and platelet aggregation were analyzed in GraphPad (v10) using two-way ANOVA with Šídák’s multiple comparisons correction, or Tukey’s multiple comparison correction as indicated. Asterisks denote statistically significant differences (*<0.05, ****<0.0001).

## Acknowledgments

We are very grateful for funding acquired from the sources mentioned below. This work benefited from access to the following Instruct-ERIC centers: the Cryo-EM facility and the Instruct Image Processing Center (I2PC) at Centro Nacional de Biotecnología (CNB-CSIC, Madrid, Spain), as well as the Institut de Biologie Structurale (IBS) and the Integrated Structural Biology Grenoble (ISBG) EM facility of Instruct-FR in Grenoble, France. Access to these Instruct-ERIC facilities was supported through Instruct-ERIC proposals. The Cryo-EM Facility CNB-CSIC and I2PC are supported by the ‘Severo Ochoa’ Programme for Centres of Excellence in R&D (CEX2023-001386-S). Cryo-EM data were collected at the CM02 beamline of ESRF under the proposal ac-21-13. The purchase of the microscope used at the CM02 CRG beamline operated by the IBS at ESRF was funded by the EquipEx+ France CryoEM project (ANR-21-ESRE-0046); we thank Guy Schoehn for the establishment of the Cryo-EM facility, and Instruct-ERIC, ESRF, and the CM02 team for providing access and support. Finally, we thank the flow cytometry facility of Sanquin Research (Amsterdam, The Netherlands) for technical support, and all individuals who volunteered to donate blood in the Sanquin Blood Bank (Amsterdam, The Netherlands).

## Data and materials availability

The structure is deposited in the Protein Data Bank (PDB) under ID 9TD2, Extended PDB ID pdb_00009TD2, and in the Electron Microscopy Data bank under ID EMD-55800.

## Author contributions

Conceptualization: JMdP, CM

Methodology: JMdP, MG, FJC, EZ, JT, CM

Validation: JMdP, MG, WS, CM

Formal analysis: JMdP, MG, WS

Investigation: JMdP, MG, WS, FvdM, FJC, EZ

Resources: FJC, EvdS, GV, JT

Visualization: JMdP, MG, WS, CM

Supervision: JMdP, CM

Writing - original draft: JMdP, CM

Writing - review & editing: JMdP, MG, WS, FvdM, FJC, EZ, EvdS, GV, JT, CM

Project administration: JMdP, CM Funding acquisition: JMdP, CM

## Abbreviations

ADMIDAS: Adjacent to MIDAS
CDR: Complementarity-Determining Regions
cryo-EM: cryo-Electron Microscopy
I-EGF: Integrin-Epidermal Growth Factor-like
Fab: Fragment Antigen-Binding region
FB: Fibrinogen
Fc: Fragment Crystallizable region
FNAIT: Fetal/Neonatal AlloImmune Thrombocytopenia
GS-FSC: Gold Standard Fourier Shell Correlation
HPA: Human Platelet Antigen
MIDAS: Metal Ion-Dependent Adhesion Site
PDB: Protein Data Bank
RMSD: Root Mean Square Deviation
SyMBS: Synergistic Metal Ion-Binding Site
TEV: Tobacco Etch Virus
PGE1: Prostaglandin E1
PSI: Plexin-Semaphorin-Integrin
PTP: Post-Transfusion Purpura

