## Supplementary Materials for "High-resolution cryo-EM structure of integrin αIIbβ3 bound to disease-causing maternal HPA-1a antibody that blocks integrin activation"

**This PDF file includes:**

Figs. S1 to S5
Table S1

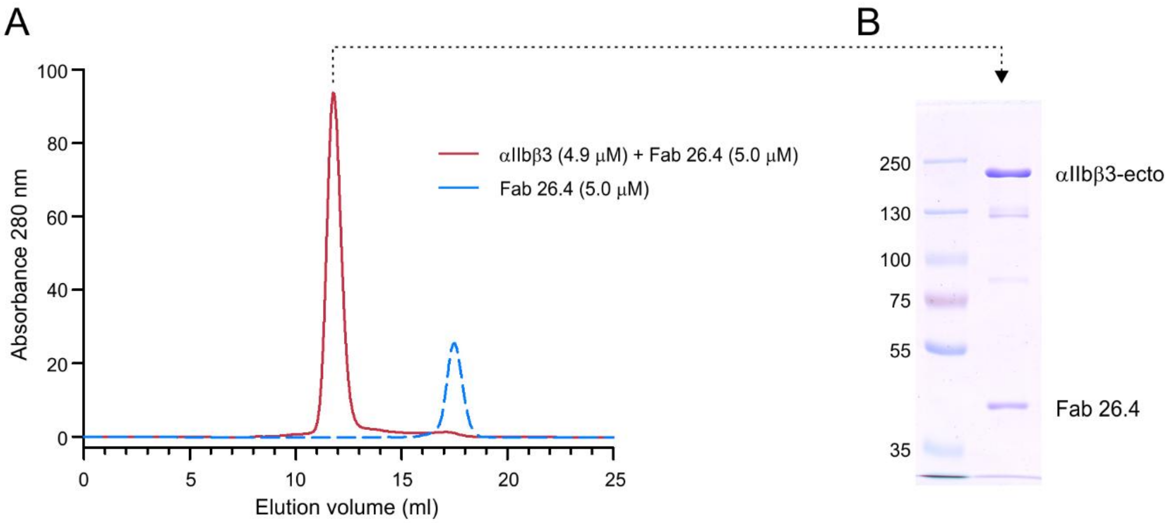


**Figure S1. Formation and purification of the Fab 26.4/αIIbβ3 complex by size exclusion chromatography.** (**A**) Elution profiles of a mixture of αIIbβ3-ecto with Fab 26.4 (*red*) or Fab 26.4 alone (*blue*) loaded at the indicated concentrations in a Superdex 200 10/300 GL column. The complex eluted as a single peak that was concentrated by ultrafiltration. In the presence of αIIbβ3, all Fab 26.4 elutes in the peak of the complex. (**B**) Analysis by SDS-PAGE (8% acrylamide gel) under non-reducing conditions of the sample in the elution peak of the complex, showing the presence of the integrin and the Fab. The faint bands at approximately 80 kDa and 125 kDa most likely correspond to a minor fraction of the β3- and αIIb-subunits, respectively, that are not linked by the disulfide bond in the coiled-coil.

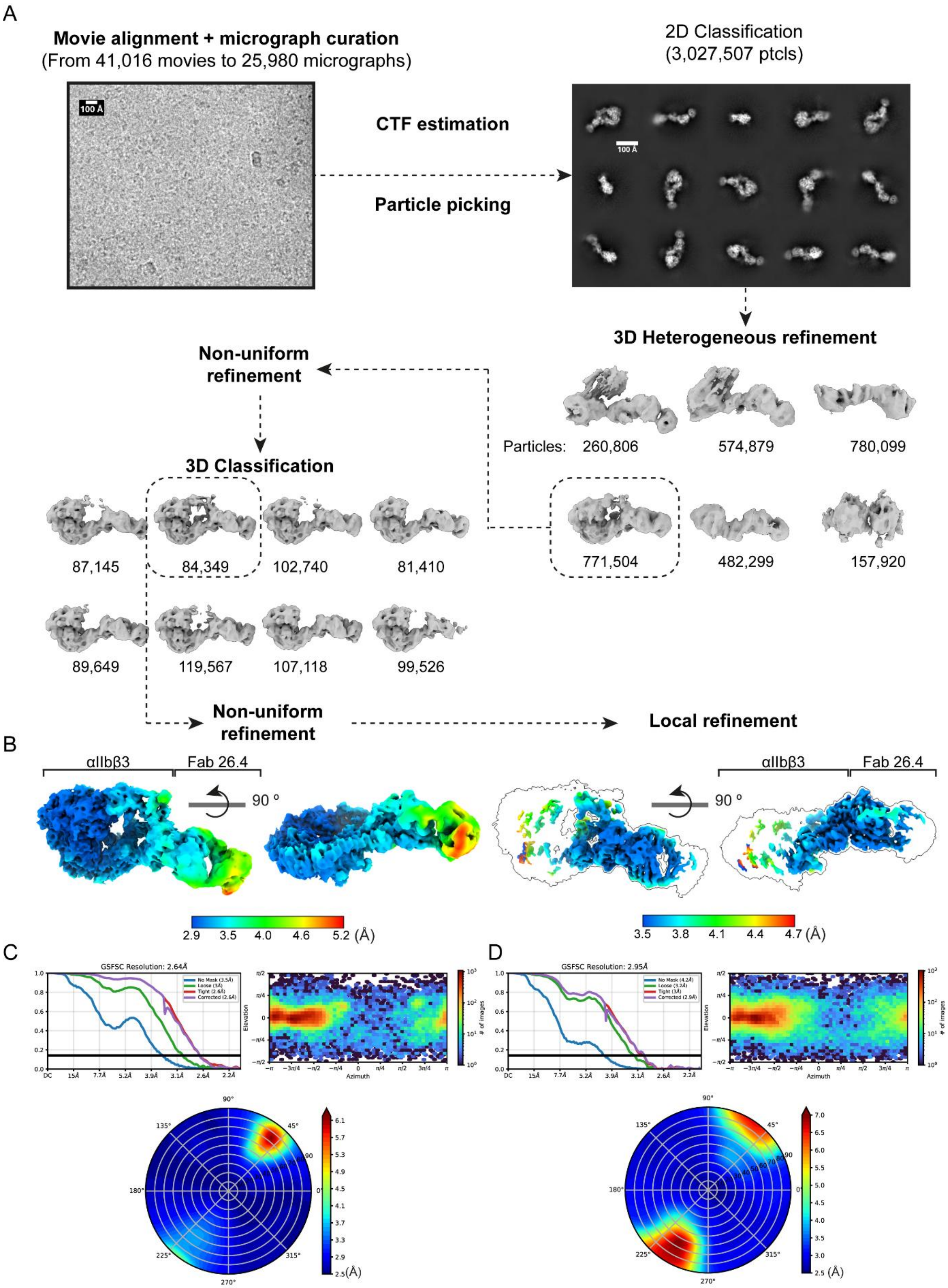
A
Movie alignment + micrograph curation
(From 41,016 movies to 25,980 micrographs)
100 Å
CTF estimation
Particle picking
2D Classification
(3,027,507 ptcls)
100 Å
3D Heterogeneous refinement
Particles: 260,806
574,879
780,099
771,504
482,299
157,920
Non-uniform refinement
3D Classification
87,145
84,349
102,740
81,410
89,649
119,567
107,118
99,526
Non-uniform refinement
Local refinement
B
αIIbβ3
Fab 26.4
90 º
2.9 3.5 4.0 4.6 5.2 (Å)
αIIbβ3
Fab 26.4
90 º
3.5 3.8 4.1 4.4 4.7 (Å)
C
GSFSC Resolution: 2.64Å
D
GSFSC Resolution: 2.95Å

**Figure S2. Cryo-EM data processing, refinement and resolution statistics.** (**A**) Flowchart of the single particle analysis pipeline. After movie alignment, micrograph curation and CTF estimation, particles were picked and 2D classified. Then, they were subjected to 3D heterogeneous refinement. The class featuring α and β subunits, together with Fab 26.4, was selected for a further round of 3D classification without recalculating angles. The class with more density was chosen for final rounds of global and local refinements. Classes selected for downstream classification/refinement are in dashed boxes. (**B**) Local resolution maps of the αIIbβ3-ecto/Fab 26.4 complex before (*left*) and after (*right*) focused refinement on the binding interface. In both panels, maps are displayed in the same two orientations. The focused-refined map is shown superimposed on a silhouette of the αIIbβ3-ecto/Fab 26.4 map. (**C, D**) Gold-Standard Fourier Shell Correlation (*left*), particle angular distribution (*right*) and directional resolution plots (*bottom*) for global and focused refinement of the αIIbβ3-ecto/Fab 26.4 complex, respectively.

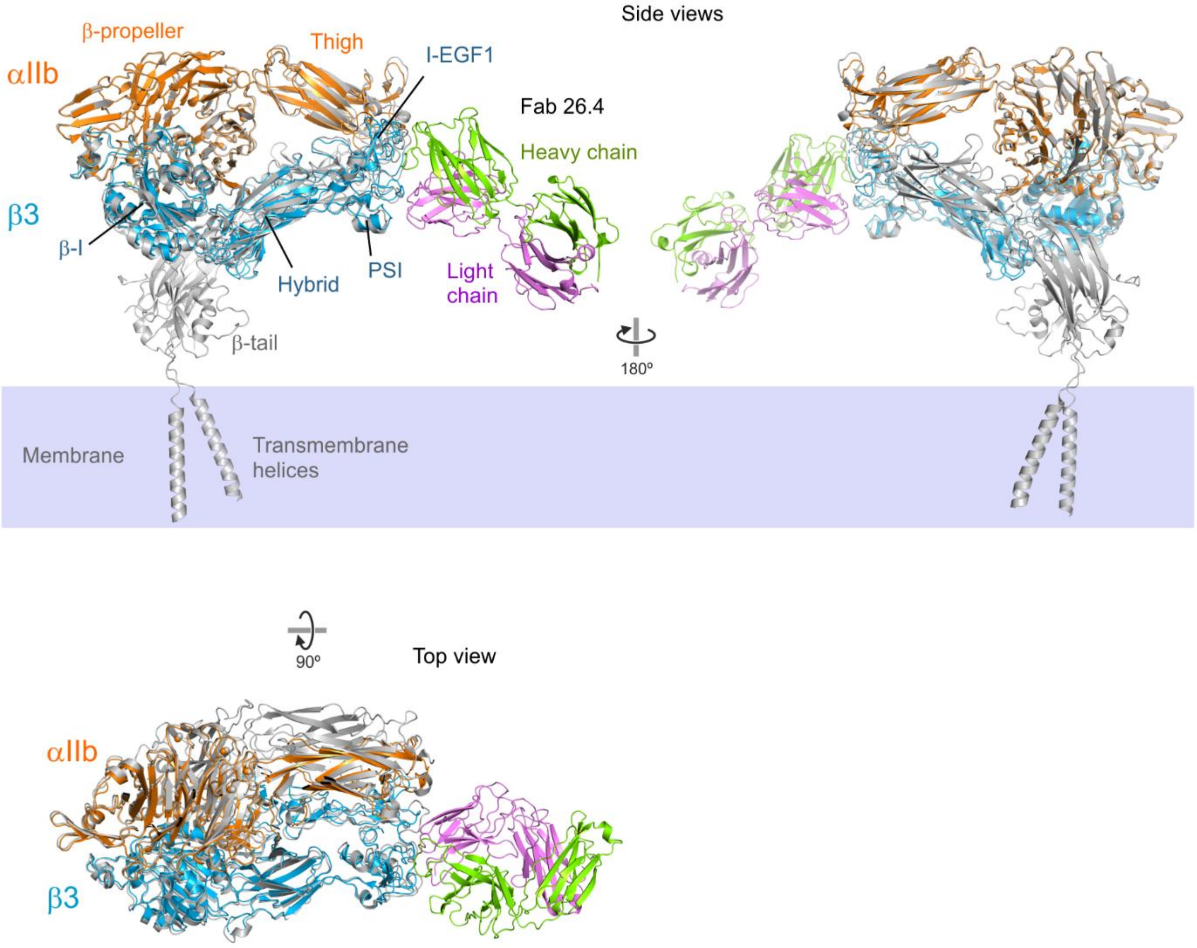


**Figure S3. Superimposition of the αIIbβ3-ecto/Fab 26.4 complex onto full-length αIIbβ3.** Comparison of the structures of the αIIbβ3-ecto/Fab 26.4 complex and full-length αIIbβ3. Ribbon representation of the cryo-EM structure of native full-length αIIbβ3 in cell-membrane nanoparticles (*grey*) (PDB: 8T2V) with the structure of the αIIbβ3-ecto/Fab 26.4 superimposed by fitting the αIIbβ3 moiety. The putative orientation of full-length αIIbβ3 with respect to the membrane is based on the position of the transmembrane helices.

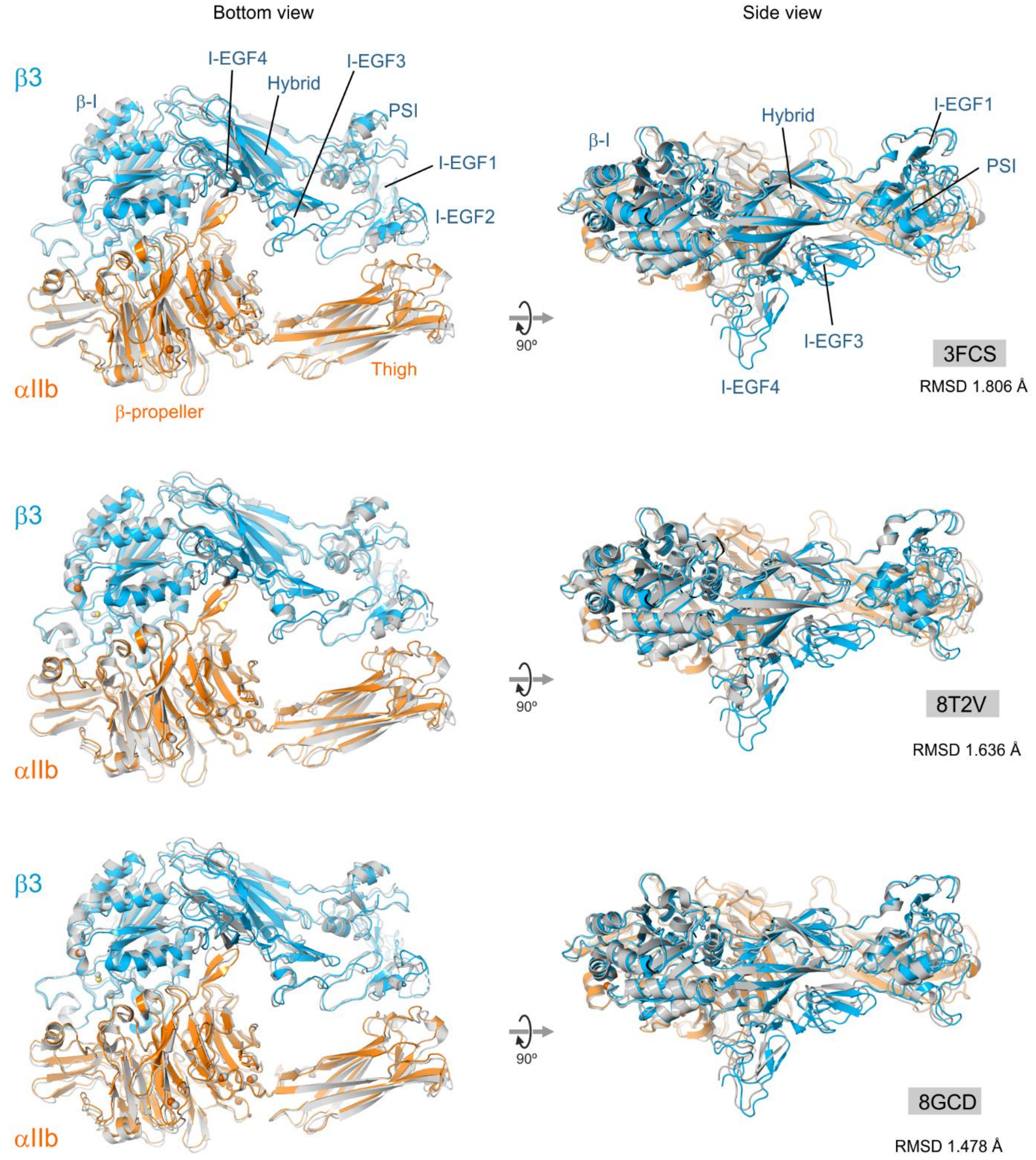


**Figure S4. Superimposition of αIIbβ3-ecto (from our αIIbβ3-ecto/Fab 26.4 complex) onto structures of αIIbβ3 in bent conformation.** Three structures of αIIbβ3 in bent conformations were pairwise superimposed onto αIIbβ3-ecto in our structure (here omitting Fab 26.4 for better comparison) by fitting the position of Cα atoms. For each superimposition two orthogonal views are shown. αIIbβ3-ecto from the complex is chain-colored (αIIb *orange* and β3 *blue*), the superimposed structure is shown in *grey*. For clarity, additional domains in the compared structures not modeled in our αIIbβ3-ecto/Fab 26.4 complex are omitted. RMSD with 3FCS was 1.806 Å for 1184 Cα atoms, with 8T2V was 1.636 Å for 1191 Cα atoms, and 1.478 Å with 8GCD for 1189 Cαs atoms.

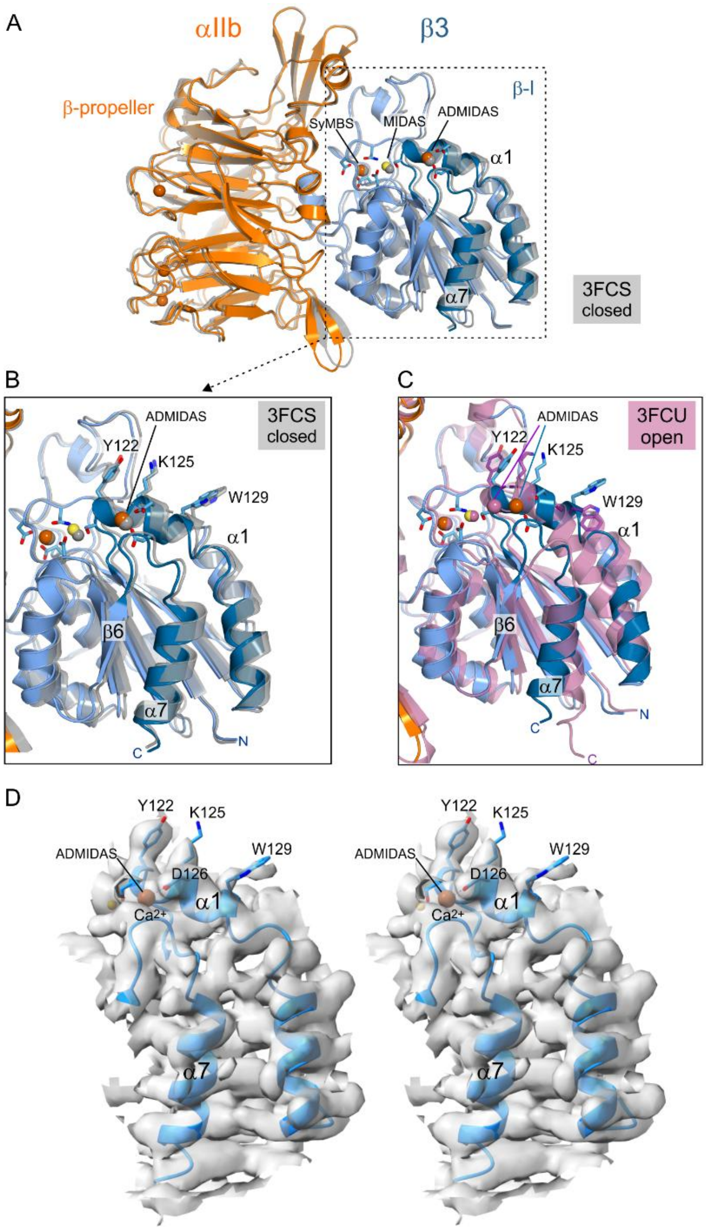


**Figure S5. The ligand-binding head region of αIIbβ3-ecto bound to Fab 26.4 is in inactive conformation.** (**A**) Ribbon representation of the head region of αIIbβ3-ecto bound to Fab 26.4 (αIIb *orange*, β3 *blue*) superimposed onto the same region of the crystal structure of αIIbβ3-ecto in the inactive conformation (*grey*) (PDB: 3FCS). (**B**) Close-up view of the region boxed in (A) corresponding to the β3 β-I-domain in the complex (*blue*) superimposed onto the inactive structure (*grey*). (**C**) For comparison, superimposition of the β3 β-I-domain in the complex (*blue*) onto the crystal structure of the αIIbβ3 headpiece in open conformation (PDB: 3FCU). There are large differences between the two structures, further supporting the notion that the β3 β-I-domain is in a closed conformation in the complex with Fab 26.4. (**D**) Stereo view of a section of the sharpened full cryo-EM map around the ADMIDAS site and helices α1 and α7 of the β3 β-I domain. A ribbon representation of this part of the structure is shown. The side chains of several residues in α1 are shown as sticks.

**Table S1. Cryo-EM data collection, refinement and validation statistics.**

| | |
|---|---|
| **Data collection & processing** | |
| Microscope | Titan Krios G4 |
| Detector | Falcon 4i |
| Magnification | 165,000x |
| Voltage (kV) | 300 |
| Electron exposure ($e^{-}$/Å$^2$) | 41.57 |
| Defocus range (µm) | -0.5 to -2.2 |
| Pixel size (Å) | 0.7 |
| Symmetry imposed | C1 |
| Image stacks (No.) | 41016 |
| Initial particle images (No.) | 6,717,934 |
| Final particle images (No.) | 84349 |
| Map resolution (Å) | 2.64 |
| FSC threshold | 0.143 |
| Map resolution range (Å) | 2.9 – 5.2 |
| Map sharpening B factor (Å$^2$) | -73.2 |
| Model resolution range | |
| dFSC model (0 / 0.143 / 0.5) (Å) | 2.6 / 2.7 / 3.3 |
| **Model refinement & validation** | |
| EMDB code: | |
| PDB code: | |
| Initial model used (PDB code) | 3FCS |
| Model composition | |
| Non-hydrogen atoms | 12766 |
| Protein residues | 1643 |
| Glycans | 16 |
| Waters | 9 |
| $Ca^{2+}$ | 6 |
| $Mg^{2+}$ | 1 |
| B factors (Å$^2$) min/max/mean | |
| Protein | 43.5 / 415.8 / 147.5 |
| Glycans | 63.0 / 261.4 / 173.1 |
| Waters | 77.7 / 104.7 / 89.0 |
| $Ca^{2+}$ | 70.3 / 124.6 / 101.6 |
| $Mg^{2+}$ | - / - / 92.9 |
| R.m.s. deviations | |
| Bond lengths (Å) | 0.004 |
| Bond angles (°) | 0.919 |
| Ramachandran plot | |
| Favored (%) | 95.46 |
| Allowed (%) | 4.35 |
| Disallowed (%) | 0.18 |
| Validation | |
| MolProbity score | 1.80 |
| Clashscore | 9.03 |
| Poor rotamers (%) | 0 |
| Map-model metrics | |
| CC(box) | 0.86 |
| CC(mask) | 0.77 |
| Average Q-score | 0.368 |
